# Dysfunctional oscillatory bursting patterns underlie working memory deficits in adolescents with ADHD

**DOI:** 10.1101/2024.12.09.627520

**Authors:** Brian C. Kavanaugh, Megan M. Vigne, Isaiah Gamble, Christopher Legere, Gian DePamphilis, W. Luke Acuff, Eric Tirrell, Noah Vaughan, Ryan Thorpe, Anthony Spirito, Stephanie R. Jones, Linda L. Carpenter

**Author notes:** Corresponding Author: Brian Kavanaugh, PsyD, ABPP, 1011 Veterans Memorial Parkway, East Providence, RI 02915.

## Abstract

Identifying neural markers of clinical symptom fluctuations is prerequisite to developing more precise brain-targeted treatments in psychiatry. We have recently shown that working memory (WM) in healthy adults is dependent on the rise and fall interplay between alpha/beta and gamma bursts within frontoparietal regions, and that deviations in these patterns lead to WM performance errors. However, it is not known whether such bursting deviations underlie clinically relevant WM-related symptoms or clinical status in individuals with WM deficits. In adolescents (n=27) with attention deficit hyperactivity disorder (ADHD), we investigated WM-related dynamics between alpha/beta and gamma bursts in relation to clinical status fluctuations. Participants repeatedly completed a visual Sternberg spatial working memory task during EEG recording as part of their participation in two research studies (n=224 person-sessions). Source localizing EEG data to each participant’s structural MRI, the rate and volume of alpha, beta, and gamma bursts were examined within the dorsolateral prefrontal cortex (DLPFC) and posterior parietal cortex (PPC). Alpha/beta and gamma bursts at the DLPFC and PPC displayed complimentary roles in WM processes. Alpha/beta bursting decreased during stimuli encoding and increased during the delay, while gamma bursting was elevated during encoding and decreased during the delay. Deviations in bursting patterns were associated with WM errors and clinical symptoms. We conclude that dysfunctional alpha/beta and gamma burst dynamics within the frontoparietal region underlie both intra-individual WM performance and WM symptom fluctuations in adolescents with ADHD. Such burst dynamics reflect a novel target and biomarker for WM-related treatment development.

## INTRODUCTION

Attention deficit hyperactivity disorder (ADHD) is a common neurodevelopmental disorder that affects approximately 10% of youth and is characterized by a persistent pattern of inattention and/or hyperactivity-impulsivity ^1,2^. Given the significant heterogeneity in the clinical presentation of ADHD, and more broadly across all neuropsychiatric disorders, empirical investigations often focus on a transdiagnostic construct or endophenotype to better understand disease pathology and clinical features ^3,4^. Working memory (WM) is a foundational component of executive functions that reflects the process of holding information ‘in mind’ to execute goal-directed behaviors ^5^. WM is one useful neurocognitive endophenotype for investigating ADHD as large effect size deficits are found in ADHD cohorts and it is one of the strongest predictors of clinical and functional outcomes, although WM deficits do not occur in every individual with ADHD and WM deficits are not specific to ADHD ^6–8^.

Successful WM is dependent on activation of the underlying frontoparietal network. This includes the posterior parietal cortex (PPC) for the encoding of spatial/sensory aspects of WM stimuli, the dorsolateral prefrontal cortex (DLPFC) for executing cognitive control demands (e.g., filtering, categorizing), and the feedforward spatial signaling and feedback control signaling between the regions ^9–12^. One recent meta-analysis of event-related EEG findings in ADHD found that ADHD was most notably characterized by weaker alpha decreases and weaker beta increases across various task demands ^13^. Specific WM findings included a weaker alpha decrease during WM encoding (e.g., ^14^), although multiple inconsistencies were noted, and none of the included studies examined gamma activity. The heterogeneity in resting state EEG for ADHD is even more striking, suggesting there is no robust physiological biomarker for the disorder, although there is also some evidence of decreased alpha/beta activity ^15^ relative to healthy controls.

One source of variability relates to the output metrics of EEG, such as the commonly used averaged power or event-related potentials, which assume that band activity is sustained over the length of trials and that a similar pattern occurs across trials. However, utilizing trial-by-trial and non-averaged approaches, it is now known that observed average power metrics can be the summation of transient, high-power oscillatory bursts ^16^. This novel oscillatory burst approach applied to band activity investigations has shown tremendous promise in better understanding beta activity in human perception ^17^, beta activity in human cognitive control ^18^, as well as alpha/beta and gamma activity in non-human primate WM ^19,20^ and human WM ^21–23^. Specific to WM, local field potential and spike recordings in non-human primates have shown that beta and gamma bursts (but not sustained averaged activity) underlie WM dynamics within the PFC ^19,20^. Beta and gamma bursts have been found to be anti-correlated, with gamma bursting increasing and beta bursting decreasing during stimuli encoding (and during WM readout), while the opposite pattern has been found during the WM delay (i.e., gamma bursting decreased, and beta bursting increased). Support for this alpha/beta bursting pattern has been recently found in healthy adult samples using EEG and magnetoencephalography ^21,23^, and preliminary support for WM-related gamma bursting has been shown in healthy children and adolescents ^22^.

Conceptually, this “push-pull” interplay between alpha/beta and gamma bursting is thought to reflect gamma bursts carrying the sensory information and alpha/beta bursts carrying the cognitive control demands of WM storage ^24,25^. More broadly, these bursting patterns are thought to potentially reflect differing oscillatory states, with a low oscillatory state involved in internal cognitive control and a high oscillatory state involved in external sensory encoding ^26^. While this conceptual model is interesting given the vulnerability of children with ADHD to be in an environmentally distractible state compared to an internal cognitive control state, this approach has not been investigated in ADHD.

The objective of the current study was to investigate whether WM-related dynamics between alpha/beta and gamma bursting patterns were associated with clinical WM status in adolescents with ADHD. We utilized trial-by-trial burst characterization and source localization approaches applied to an MRI/EEG dataset of n = 224 participant/sessions (participant sample size n = 27) collected during a Sternberg working memory paradigm. As described below, alpha/beta and gamma bursting follow inverse patterns of rising and falling during WM demands in the DLPFC and PPC regions. WM errors and symptoms in these adolescents were related to deviations in these bursting patterns. Such findings provide novel markers of disease status and treatment targets in ADHD and other conditions with WM deficits.

## METHODS

### Participants/Design

Data was used from twenty-seven adolescents with a clinical diagnosis of ADHD and elevated parent-reported WM symptoms (i.e., > 1 SD above the normative mean) who participated in one or two ongoing studies investing the potential utility of intermittent theta burst stimulation (iTBS) in improving WM and ADHD symptoms (https://clinicaltrials.gov/study/NCT05662280 & https://clinicaltrials.gov/study/NCT05102864). Clinical and demographic data are provided in Table 1. In study NCT05662280, participants had EEG recorded during WM tasks (EEG/WM) immediately before and after an iTBS session on two separate days. In the sham-controlled, crossover clinical trial (NCT05102864), participants received a total of 20 iTBS sessions spread across two 11-day phases (6+ week washout between phases), including 10 active iTBS sessions and 10 sham iTBS sessions. EEG/WM is completed on days 1, 2, 6, and 11 of each phase. For the initial analyses, all available EEG data was analyzed across both studies (participant n = 27; participant/session n = 224; average per person = 6.8). For analyses that examined correlation to clinical symptoms, only data from the clinical trial were analyzed to track symptom change across days (participant n = 20; participant/session n = 135; average per person = 8.3).

**Table 1.** Demographic and Clinical Data (n = 27)

|  |  |
| --- | --- |
| Age, mean (SD) | 14.9 (1.7) |
| Gender, n (%) |  |
| Male | 18 (67) |
| Female | 8 (30) |
| Non-binary | 1 (4) |
| Academic Services, n (%) |  |
| IEP | 11 (41) |
| 504 Plan | 11 (41) |
| IQ, mean (SD) | 108.2 (10.2) |
| Handedness |  |
| Right | 24 (89) |
| Left | 2 (7) |
| Ambidextrous | 1 (4) |
| Parent Primary Language |  |
| English | 17 (100) |
| Parent 1 Education, n (%) |  |
| High School Diploma | 1 (4) |
| Some College | 3 (11) |
| 2-Year College Degree | 3 (11) |
| 4-Year College Degree | 4 (15) |
| Master's Degree | 14 (52) |
| Doctoral Degree | 2 (7) |
| Parent 2 Education, n (%) |  |
| Not applicable | 6 (22) |
| High School Diploma | 3 (11) |
| Some College | 2 (7) |
| 2-Year College Degree | 1 (4) |
| 4-Year College Degree | 3 (11) |
| Master's Degree | 8 (30) |
| Doctoral Degree | 1 (4) |
| Child's Race, n (%) |  |
| White | 20 (74) |
| Multi-race | 3 (12) |
| Other | 4 (15) |
| Child's Ethnicity, n (%) |  |
| Hispanic | 4 (15) |
| Non-Hispanic | 23 (85) |
| Household Income |  |
| \$26,000 to \$49,999 | 1 (4) |
| \$50,000 to \$74,999 | 6 (22) |
| \$75,000 to \$99,999 | 2 (7) |
| \$100,000 to \$149,000 | 7 (26) |
| \$150,000 or more | 11 (41) |
| Psychiatric Diagnoses, n (%) |  |
| Attention deficit hyperactivity disorder | 27 (100) |
| Anxiety disorders | 16 (59) |
| Depressive disorders | 11 (41) |
| Eating disorders | 2 (7) |
| Obsessive-compulsive disorder | 1 (7) |
| Oppositional defiant disorder | 2 (7) |
| Learning disorders | 1 (4) |
| Autism spectrum disorder | 1 (4) |
| Disruptive mood dysregulation disorder | 1 (4) |
| Mood disorders | 2 (7) |
| Psychiatric Medications, n (%) |  |
| Antidepressant | 15 (56) |
| Antihistamine | 3 (11) |
| Antipsychotic | 1 (4) |
| Stimulant | 20 (74) |
| Non-stimulant | 8 (31) |

### Working Memory Task

All participants completed a computerized, visuospatial Sternberg working memory paradigm (**Figure 1.A**) ^14,27^. During encoding (2 seconds), yellow dots (either 1, 3, 5, or 7) were presented on the screen, while in the probe condition, a single green dot was presented (up to 3 seconds). For consistency across datasets, only trials with 3, 5, or 7 dots were analyzed. See Supplement for additional details.

**Figure 1.**
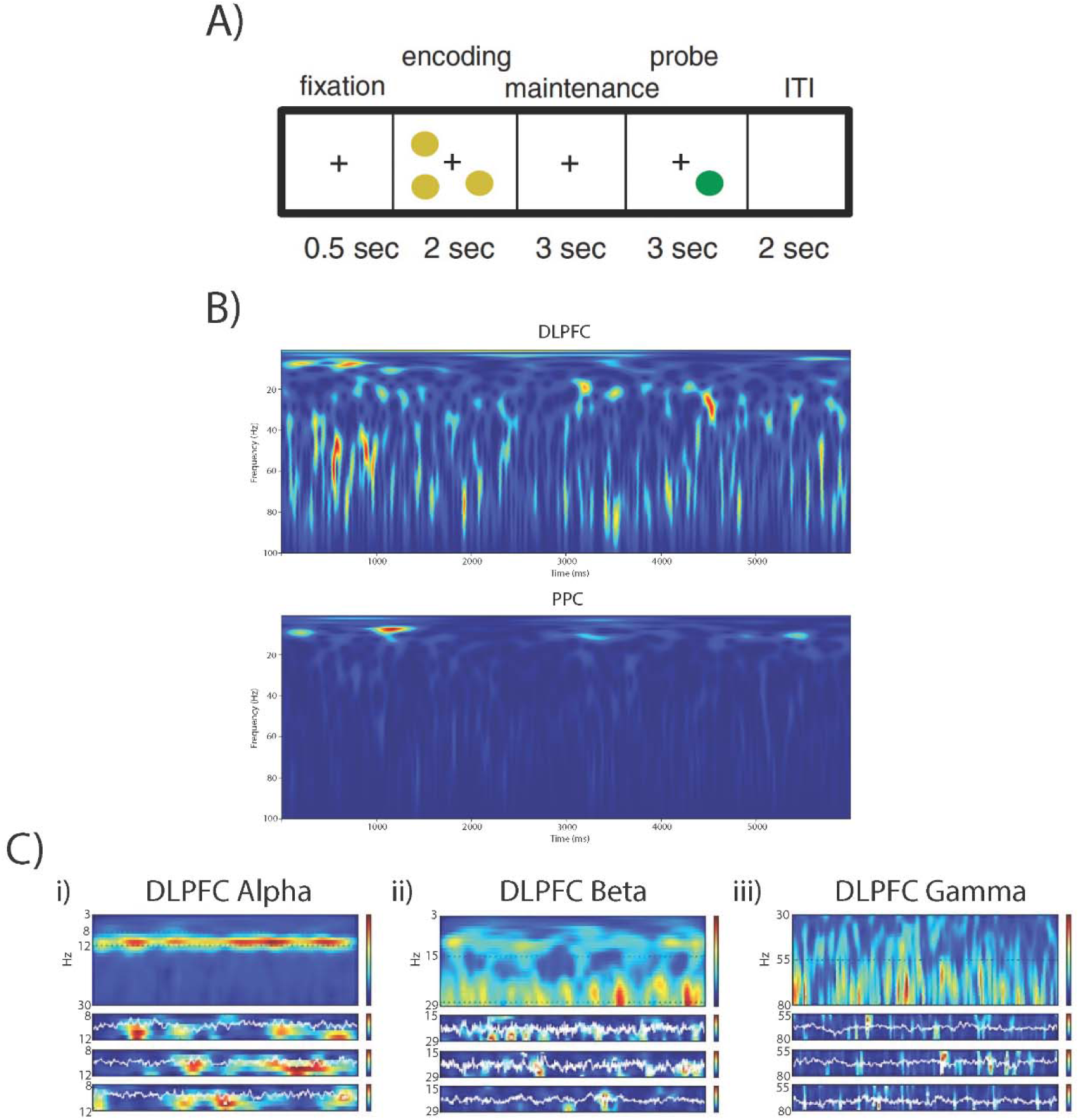
Initial spectrogram and sample burst raw data. A) The experimental paradigm. Yellow dots (3, 5, or 7) were used as encoding stimuli, with one green dot as the stimuli during the probe. As previously described by Lenartowicz et al. (2014). B) Spectrogram across bands at DLPFC and PPC. C) Sample averaged oscillatory power during individual recordings, decomposed into distinct alpha (left), beta (center), and gamma (right) bursts across three randomly sampled trials.

### Clinical Variables

Parents completed the National Institute for Children’s Health Quality (NICHQ) Vanderbilt Assessment Scale for ADHD^28^ as well as the working memory subscale of the Behavior Rating Inventory of Executive Function – Second Edition (BRIEF-2)^29^. The ADHD total score sum from the NICHQ and the total sum score of the BRIEF-2 working memory subscale were completed by the parent electronically during each study visit. Stimulant and non-stimulant medication status were parent-reported at the intake session.

### EEG Recording & Preprocessing

In both studies, the same procedural format was followed for each day that there was an EEG recording. EEG recording was obtained during the Sternberg working memory task with a 64-electrode montage integrated in a neoprene cap (Waveguard by ANT Neuro). Pre-processing was conducted using the EEGLAB toolbox for MATLAB. See Supplement for additional details.

Data were epoched around the onset of the fixation prior to trial onset, with a baseline correction of −.5 to 0 (reflecting the fixation onset) to capture baseline/fixation (500 milliseconds), encoding (2000 milliseconds), maintenance (3000 milliseconds), and the initial portion of the probe (500 milliseconds). When examining discrete WM stages, 500 milliseconds EEG collected during the fixation cross was used as baseline. Specifically, the middle second of encoding was extracted (i.e., 0.5-1.5 from the available 0 – 2.0 seconds). For the delay, the middle two seconds were extracted (i.e., 0.5-2.5 from the available 0 - 3.0 seconds). The first 500 milliseconds of the probe were extracted to capture EEG data prior to motor response.

### Source Localization

Estimated source localization of DLPFC (rostral-dorsal area of region 39) and PPC (ventral area of region 9/46) regions of interest was conducted through Brainstorm, an open-source EEG/MEG data processing application using the Brainnetome Atlas ^30^. The left and right regions were averaged for a single DLPFC and a single PPC. See Supplement for additional details.

### EEG Spectral Analysis

Transient high-power “events” were detected and characterized using the SpectralEvents Toolbox (https://github.com/jonescompneurolab/SpectralEvents). A spectral event was defined as any local maximum in the time-frequency response (TFR) above a power threshold within a user-defined band of interest. To be consistent with prior studies using similar methods, we used findMethod=1^17^, which is agnostic to event overlap. The event threshold was set at 6x the median power (i.e., 6 factors-of-the-median [FOM]) across time and epochs for each frequency bin of the TFR ^17,31,32^. Events were examined within the alpha (8-12 Hz), beta (15-29 Hz) and gamma (55-80 Hz) bands based on our recent work in healthy young adults (^33^; **Figure 1**). Each spectral event was characterized by its peak time/frequency within each trial, along with the event’s peak power, duration, and frequency span (f-span). Analysis was conducted on a subject-by-subject basis. *Event rate* was calculated by counting the number of events within each trial. *Event volume* was calculated as the mean of z-scores for event power, event duration, and event frequency span. *Event power* was calculated as the normalized FOM power value at each event maximum. The *Event duration* and *frequency span* (F-span) were calculated from the boundaries of the region containing power values greater than half the local maxima power, as the full-width-at-half-maximum in the time and frequency domain, respectively. Events with characteristics greater than three standard deviations from the mean were removed. Z-scores of event characteristics are reported.

### Statistical Analyses

Statistical analyses were conducted in Matlab-2022a and GraphPad Prism 10. One- and two-way repeated measures analysis of variance (RM-ANOVA) were utilized for most analyses, although if missing data was present, a mixed-effects model was alternatively conducted. For the first set of initial analyses, the primary effect of time (i.e., fixation, encoding, delay, probe) was examined as indicated. Analyses examining differences between region or accuracy utilized the time x accuracy/region metric in the model. Pairwise comparisons after models were corrected with Sidak correction. Unless otherwise indicated, correct trials were utilized. Bonferroni correction to account for two burst features (rate and volume) and three bands (alpha, beta, gamma) was applied to all initial analyses, including RM-ANOVA and mixed models (i.e., raw p-value * 6 = corrected p-value). Based on these initial results, we examined whether the difference between alpha/beta and gamma bursting was related to task accuracy (i.e., gamma - alpha/beta contrast). We first calculated the difference between gamma bursting z-score and the average of the alpha + beta bursting rate z-score ((alpha z-score + beta z-score)/2). We then examined the differences in this alpha/beta – gamma contrast score between incorrect and correct trials across time at the DLPFC and PPC for burst rate only. This gamma-alpha/beta contrast (G-ABC) score was utilized for all analyses related to clinical status. For the clinical status analyses (subsample participant/sessions = 135), Bonferroni correction is applied to account for examining both DLPFC and PPC. Raw burst scores converted to z-scores were utilized for analyses unless indicated otherwise. Bonferroni corrected p-values are reported, with statistical significance set at p < .05.

## RESULTS

### Alpha/beta and gamma bursting have partially inverse patterns across WM demands

#### PPC

<u>Alpha/Beta</u>: Alpha and beta bursting followed a similar pattern across WM stages, decreasing in rate and volume of bursts from fixation to encoding, then increasing from encoding to the delay. There was also evidence of alpha increasing and beta decreasing activity from the delay to the probe (Alpha burst rate: F = 9.0, p < .0001; Alpha burst volume: F = 14.4, p < .0001; Beta burst rate: F = 43.1, p < .0001; Beta burst volume: F = 18.3, p < .0001). <u>Gamma</u>: In concert with these changes, the rate and volume of gamma bursts decreased from encoding to the delay (while alpha/beta was increasing) and further decreased from the delay to the probe (while alpha/beta was also decreasing; **Figure 2.B**; Gamma burst rate: F = 24.1, p < .0001; Gamma burst volume: F = 16.3, p < .0001).

**Figure 2.**
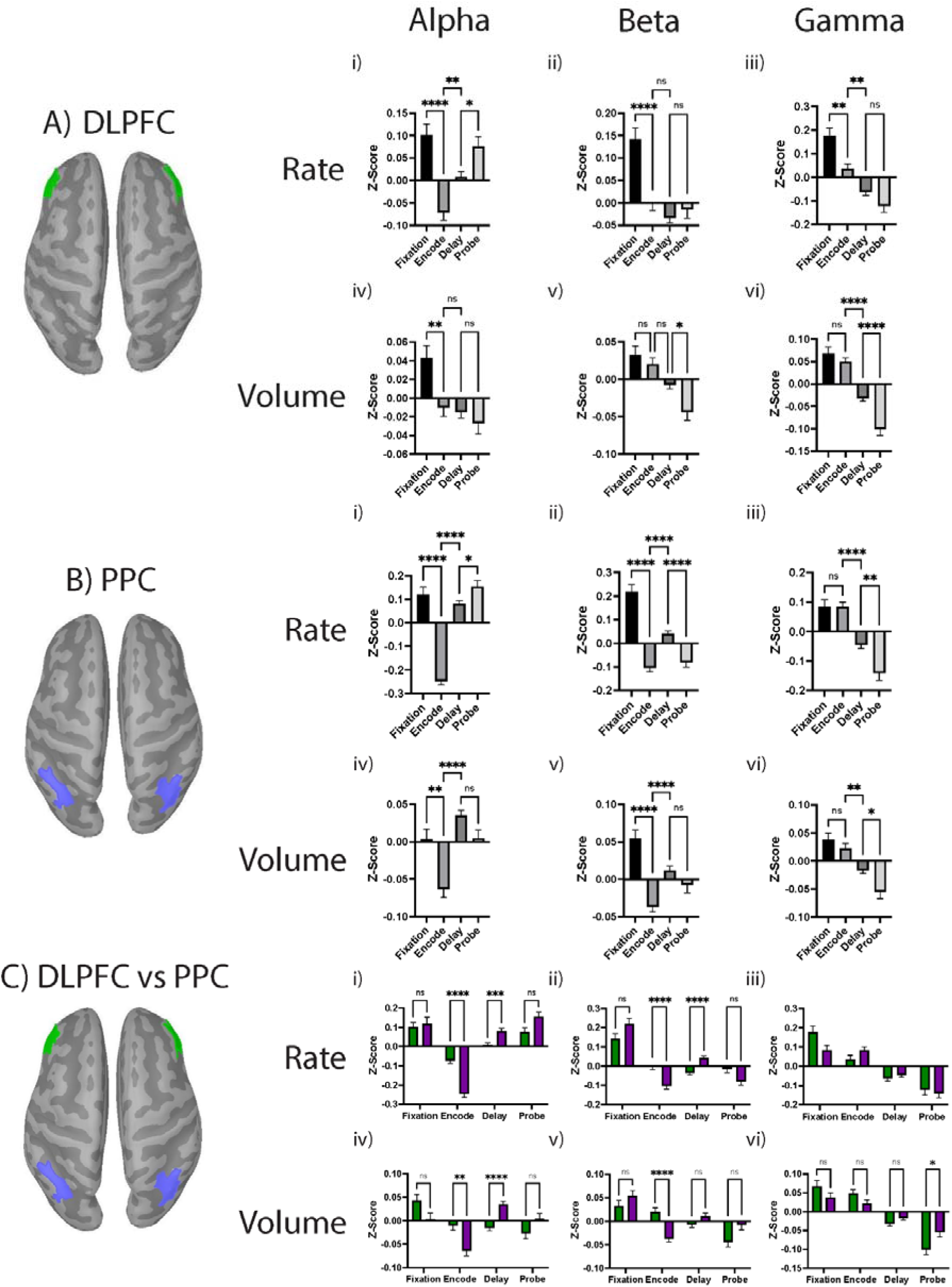
Alpha/beta and gamma bursting have partially inverse patterns across WM demands. A) Within the DLPFC, the rate of alpha/beta bursting decreased from fixation to encoding and beta additionally increased from encoding to the delay. The rate and volume of gamma bursting decreased from encoding to delay and from delay to probe. B) Within the PPC, the rate and volume of alpha/beta bursting decreased from fixation to encoding and increased from encoding to the delay. There was also some evidence of decreasing from delay to probe. The rate of gamma bursting decreased from encoding to the delay and from the delay to the probe. C) The dynamic variation in alpha/beta bursting was more pronounced in the PPC compared to the DLPFC. Primary models were Bonferroni corrected, with pairwise comparisons (Sidak correction) only shown if model was p < .05.

#### DLPFC vs PPC

<u>Alpha/Beta</u>: Compared to the PPC, there was less dynamic variation (i.e., less pronounced variation in bursting) of alpha and beta bursting in the DLPFC, seen particularly in the rate and volume of alpha bursts and in the rate of beta bursts during the encoding and delay stages (**Figure 2.C**; PPC vs DLPC: Alpha burst rate: F = 12.3, p < .0001; Alpha burst volume: F = 11.7, p < .0001; Beta burst rate: F = 9.4, p < .0001; Beta burst volume: F = 20.5, p < .0001). <u>Gamma</u>: There were minimal differences in gamma bursting between DLPFC and PPC (Gamma burst rate: F = 2.9, p > .05; Gamma burst volume: F = 58.9, p < .0001).

#### DLPFC

<u>Alpha/Beta</u>: Within the DLPFC, and similar to PPC findings, the rate of alpha bursting decreased from fixation to encoding and then increased from encoding to the delay and from the delay to the probe (**Figure 2.A**; Alpha burst rate: F = 13.2, p = .0001), but only a decrease from fixation to encoding was observed for bursting volume (Alpha burst volume: F = 8.3, p < .0001). Similarly, the rate of beta bursting in the DLPFC decreased from fixation to encoding (Beta burst rate: F = 15.2, p < .0001), but no further increase was observed from encoding to the delay. A decrease from delay to probe was observed for beta bursting volume (Beta burst volume: F = 12.8, p = < .0001). <u>Gamma:</u> Both the rate and volume of gamma bursting steadily decreased from fixation to the probe (Gamma burst rate: F = 21.3, p < .0001; Gamma burst volume: F = 44.2, p < .0001).

### Incorrect responses were associated with reduced dynamic variation in bursting patterns

Within the DLPFC, correct trials (compared to incorrect trials), had higher beta burst rate and higher alpha burst volume during fixation, higher gamma burst volume during encoding, and lower beta burst rate during the delay (**Figure 3.A**; Beta burst rate: F = 6.1, p = .002; Alpha burst volume: F = 4.9, p = .02; Gamma burst volume: F = 4.6, p = .02). Within the PPC (**Figure 3.B**), correct trials had a higher beta burst rate and volume during the fixation, higher gamma burst rate and lower alpha burst rate during encoding, and lower gamma burst rate during the probe (Alpha burst rate: F = 5.6, p = .005; Beta burst rate: F = 12.3, p < .0001; Gamma burst rate: F = 6.1, p = .002; Beta burst volume: F = 4.6, p = .02). <u>Gamma - Alpha/Beta Contrast</u>: In the PPC, but not the DLPFC (p > .05), correct trials (compared to incorrect trials) were characterized by a larger contrast between gamma and alpha/beta burst rate across time (**Figure 3.C**; F = 6.9, p = .018).

**Figure 3.**
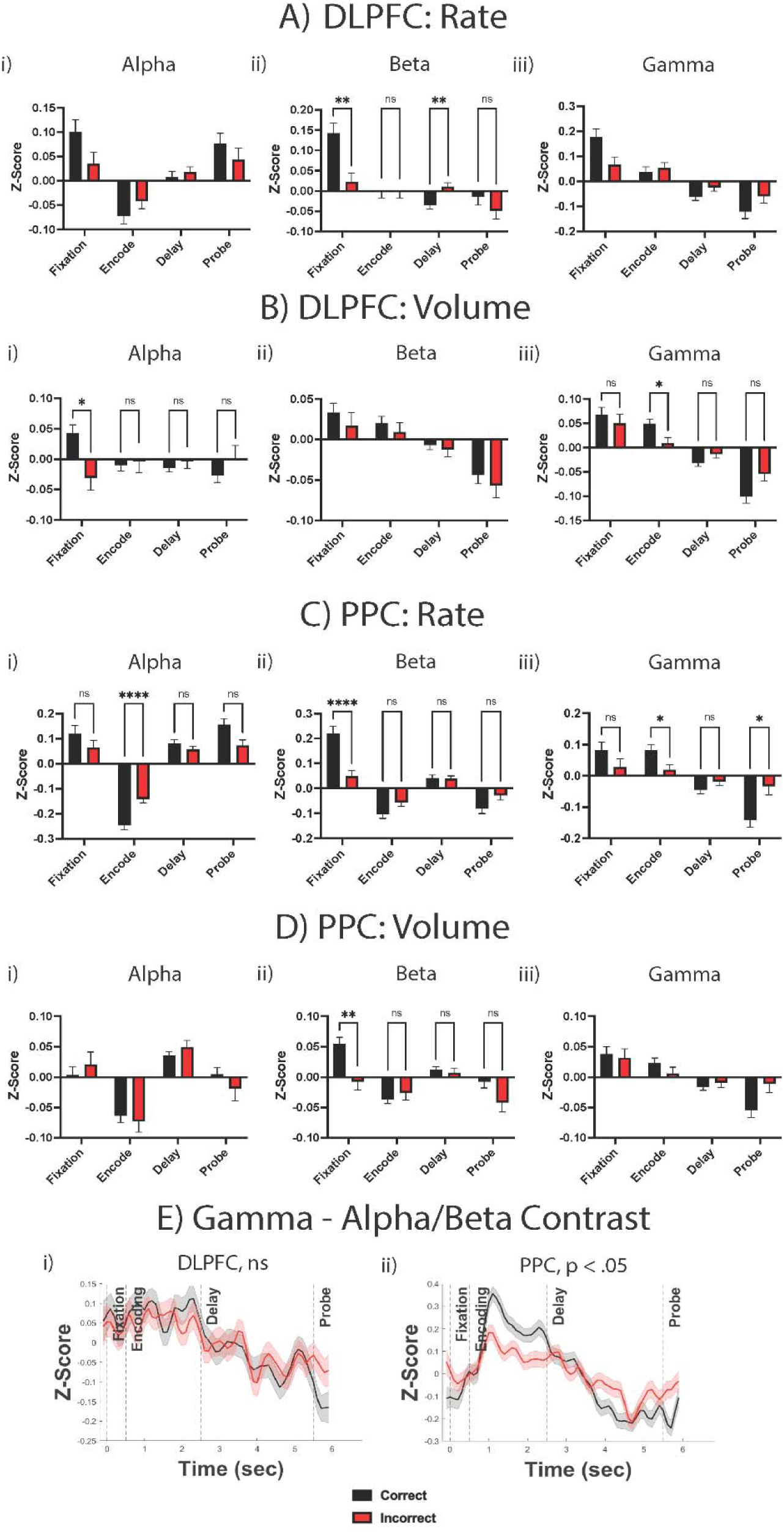
Incorrect responses were associated with reduced dynamic variation bursting patterns. Bursting pattern dynamic differences between correct and incorrect trials were examined within the DLPFC/PPC regions and for burst volume and rate. Primary models were Bonferroni corrected, with pairwise comparisons (Sidak correction) only shown if model was p < .05.

### Aberrant bursting dynamics are related to clinical status

In the DLPFC, G-ABC was correlated with visit number, in that as visits increased, G-ABC increased during encoding (i.e., gamma increased), decreased during the delay (i.e., alpha/beta increased), and decreased during the probe (i.e., alpha/beta increased; **Figure 4.A & 4.J**). In the PPC, WM performance was correlated with G-ABC during fixation and the probe, in that higher WM performance was associated with lower G-ABC during the fixation (i.e., alpha/beta higher) and higher G-ABC during the probe (i.e., gamma higher; **Figure 4.A & 4.B**). G-ABC during the fixation visually tracked with WM performance fluctuations across study visits (**Figure 4.C**). ROC curve results indicated that G-ABC was able to detect low WM performance (i.e., < 25^th^ percentile of sample; AUC = .70, p = .004; **Figure 4.D**) and the low WM performing group had higher G-ABC during the fixation than the rest of the sample (> 25^th^ percentile; t = 3.2 [108], p = .004; **Figure 4.E**).

**Figure 4.**
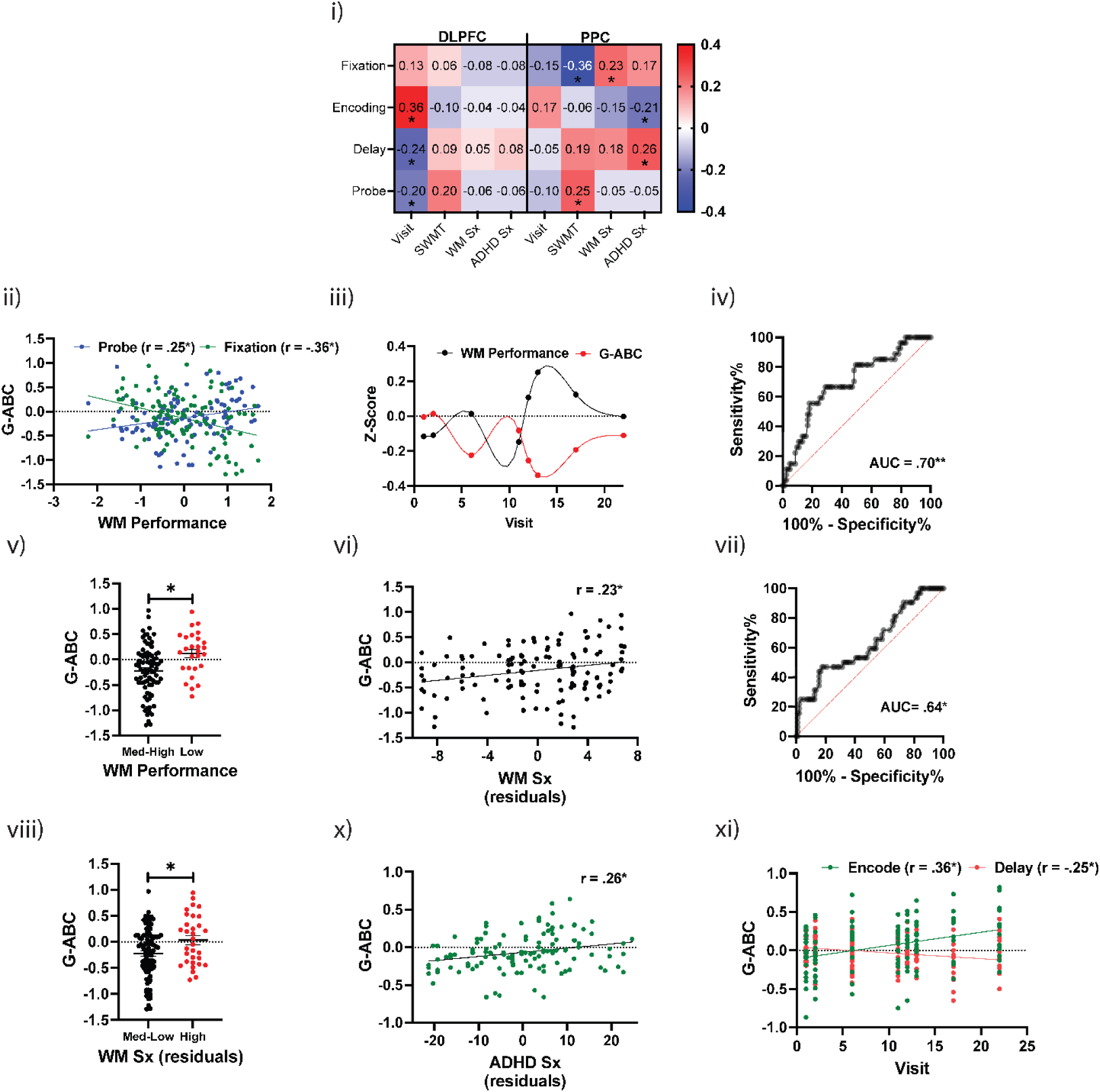
Dysfunctional bursting dynamics are related to clinical status. i) Heatmap of correlation between G-ABC during different WM stages and relevant clinical variables. ii) Association between WM task performance and G-ABC within the PPC. iii) Sample average WM performance and G-ABC during fixation in the PPC with spline curve line across study visits. iv) ROC curve results of G-ABC during fixation in the PPC in detecting low (i.e., < 25^th^ percentile of sample) WM task performance. v) Difference in G-ABC during fixation in the PPC between low WM task performance and rest of the sample (i.e., > 25^th^ percentile). vi) Correlation between G-ABC during fixation in the PPC and parent-reported WM symptoms (BRIEF-2 WM Subscale; age regressed out). vii) ROC curve results of G-ABC during fixation in the PPC in detecting high (i.e., > 75^th^ percentile of sample) WM symptoms. viii) Difference in G-ABC during fixation in the PPC between high WM symptoms and rest of the sample (i.e., < 75^th^ percentile). viv) Correlation between G-ABC during fixation in the PPC and parent-reported ADHD symptoms (Vanderbilt; age regressed out). x) Correlation between G-ABC in the PPC and study visit number. Bonferroni corrected for two regions (DLPFC/PPC).

Higher parent-reported and age-controlled WM symptoms was associated with higher G-ABC during the fixation in the PPC (**Figure 4.A & 4.F**). ROC curve results indicate that G-ABC was able to detect high WM symptoms (i.e., > 75^th^ percentile of sample; AUC = .64, p = .04) and the high WM symptoms group had higher G-ABC during the fixation than the rest of the sample (i.e., < 75^th^ percentile; t = 2.7 [126], p = .02). Finally, higher parent-reported and age-controlled ADHD symptoms were associated with lower G-ABC during encoding and higher G-ABC during the delay in the PPC (**Figure 4.A & 4.J**). Stimulant prescription was associated with higher G-ABC during the probe in the PPC (r = .23, p = .02) and non-stimulant prescription was associated with lower G-ABC during the delay in the PPC (r = −.21, p = .02).

## DISCUSSION

Utilizing novel, trial-by-trial burst characterization and source localization approaches applied to an MRI/EEG-WM dataset of adolescent ADHD, we demonstrated that dysfunctional gamma and alpha/beta bursting were related to WM errors and WM symptoms, potentially reflecting different oscillatory states underlying ADHD-related features. Such findings may provide novel markers of disease status and inform treatment targets for ADHD and other conditions characterized by WM deficits.

Our analyses of EEG data from an ADHD sample revealed a temporal burst pattern that reflected the pattern observed in healthy adult and non-human primate findings ^19–21^. Specifically, alpha/beta bursting decreased during stimuli encoding and increased during the WM delay, while gamma bursting was elevated during stimuli encoding and decreased during the delay. This type of alpha/beta – gamma pattern was discovered within single PFC neurons in non-human primates ^19,20^. A reciprocal pattern of alpha/beta activity during WM was reported in healthy adults ^21,34^ and we recently built off these findings in healthy young adults ^33^. Successful navigation of WM demands is known to involve an increase in alpha/beta bursting during the task initiation, deactivation of alpha/beta bursting during stimuli presentation to allow for gamma bursting and perceptual encoding, followed by increase in alpha/beta bursting to hold control of the WM information during the delay ^24^.Our current results show that this alpha/beta – gamma bursting pattern in adult and primate WM is also observed in adolescent ADHD.

We found that deviations in the alpha/beta versus gamma bursting dynamics were related to WM task errors. WM errors (i.e., correct compared to incorrect trials) were associated with a) dampened alpha and beta bursting during the fixation, b) heightened alpha bursting and dampened gamma bursting during stimuli encoding, as well as c) heightened beta bursting during the delay and heighted gamma bursting during the probe. This pattern of errors reflected an overall reduced dynamic variation in the rise-and-fall bursting pattern, in that errors were related to insufficient bursting when bursting is expected to be high (e.g., alpha/beta during fixation) and excessive bursting when bursting is expected to be low (e.g., alpha during encoding). While this is the first study to identify WM bursting patterns in either a clinical or pediatric sample, it does replicate our recent healthy young adult findings ^33^ and is broadly consistent with prior findings on the association between beta activity and WM performance in non-human primates ^35^, adults with Parkinson’s disease ^36^, and healthy adults ^37^.

This mirrored pattern between alpha/beta and gamma bursting was most succinctly captured with the gamma – alpha/beta bursting contrast score (G-ABC). In the PPC, but not DLPFC, WM errors were associated with diminished gamma predominance (i.e., lower gamma > alpha/beta bursting) during stimuli encoding, and diminished alpha/beta predominance (i.e., lower alpha/beta > gamma bursting) during the fixation, delay, and probe periods. This appears to reflect an intermittent switching during different WM demands between an alpha/beta predominance, or a low oscillatory state responsible for cognitive control processes, and a gamma predominance, or a high oscillatory state responsible for sensory perception processes, with WM errors associated with inefficient switching or balancing between these oscillatory states^24,26^. This is not inconsistent with prior functional MRI findings on anticorrelated or mirrored neural activity between functional networks ^38,39^, and suggests that EEG-measured temporal dynamics have promise in identifying WM-relevant neural network dynamics.

We then found that G-ABC in the PPC, but not the DLPFC, was tightly correlated with clinical status during a clinical trial. Lower G-ABC during the fixation (i.e., higher alpha/beta bursting) was associated with both better WM performance and lower parent-reported WM symptoms, while higher G-ABC during the probe (i.e., higher gamma bursting) was associated with better WM performance. Elevated ADHD symptoms were additionally associated with a higher level of alpha/beta bursting during encoding and lower level of alpha/beta bursting during the WM delay. This is highly consistent with meta-analytic findings from ADHD studies (using averaged power, not bursts) that show excessive alpha activity (i.e., particularly during WM encoding) and insufficient beta activity across task demands ^13,14^. Findings on G-ABC during the fixation are not inconsistent with prior findings on lowered alpha/beta activity during resting state in ADHD ^15^, and suggests that WM-related deficits may not solely relate to neural dynamics *during* WM demands, but rather neural dynamics *prior to* WM demands. From a theoretical perspective, our current findings suggest that WM deficits (and related ADHD symptoms) may arise from a dysfunctional allocation or switching between low (i.e., cognitive control) versus high (i.e., sensory encoding) oscillatory states, in that cognitive control may be incorrectly maximized when perception is needed, and/or perception is incorrectly maximized when cognitive control is needed.

Critically, G-ABC during the fixation was able to detect low WM performance and high WM symptoms, and negatively tracked with WM performance variability across days in the trial (i.e., when WM increased, G-ABC decreased). This G-ABC metric may reflect a feasible, EEG-based biomarker of disease status when tracking and optimizing target engagement or treatment response to novel neurotherapeutics in psychiatry ^40^. Targeting these dysfunctional bursting patterns is certainly possible with non-invasive brain stimulation protocols ^41^. Although not yet commercially available, closed loop or adaptive stimulation systems which would allow for the specific targeting of these bursting patterns during WM demands are particularly promising. For example, specifically enhancing alpha/beta or diminishing gamma during the fixation and during the WM delay or enhancing gamma and diminishing alpha/beta during the WM encoding phase could potentially enhance performance or relieve clinical symptoms. Though in a less specific manner, pharmacotherapies and cognitive/behavioral therapies may also be able to target these burst dynamics to improve clinical symptoms.

We focused our investigation on MRI/EEG-source-localized DLPFC and PPC regions given the established role of the frontoparietal network in human WM ^9–12^. Interestingly, while notable findings were seen in both regions, overall, the most prominent findings were within the PPC. The PPC was characterized by larger dynamic variation in bursting patterns, particularly in alpha/beta bands, and more consistent associations between bursting and WM errors and symptoms. Alternatively, we found that G-ABC normalized with increased task practice in the DLPFC, but not the PPC, in that G-ABC increased with visits during encoding and decreased with visits during the delay and probe. While the PPC bursting was more consistently associated clinical status, this learning effect suggests that the DLPFC may be more modifiable or neuroplastic, and potentially a more promising WM treatment target than the PPC. This is the first study to our knowledge that has utilized individual MRIs to calculate EEG-measured bursting characteristics in humans, and suggests that this may reflect a novel, feasible approach to measuring frontoparietal bursting dynamics in human WM. More broadly, shifting from averaged power metrics to more precise oscillatory burst measurements may further advance our understanding of neurocognition and clinical conditions.

This study has several limitations. First, while the data included 224 participant/sessions, there were only 27 unique adolescents with ADHD represented in the sample. The use of repeated intraindividual measures has utility, as we showed how burst activity tracked with WM status when subjects was tested on multiple occasions over time. However, such an approach is less optimal than a larger sample of individuals with a single time point assessment. We used each person’s structural MRI to source-localize bursting patterns to the DLPFC and PPC regions, and this method has inherent limitations in maximizing spatial resolution, compared to functional MRI. Such limitations may be more relevant for pediatric studies because the Brainnetome Atlas is based on adult brains. As there was no healthy control group, we cannot draw conclusions about abnormal neural bursting that is unique to ADHD as a disease or disorder, but rather we are limited to conclusions linked to variability in WM/ADHD symptom severity among adolescents with ADHD who are 12-18 years old.

## SUPPLEMENT

### Working Memory Task

A cross was presented in the middle of the screen for the fixation (.5 seconds) and maintenance conditions (3 seconds). During encoding (2 seconds), yellow dots (either 1, 3, 5, or 7) were presented on the screen, while in the probe condition, a single green dot was presented (up to 3 seconds). The individual was asked to determine if the green dot is in the same place as any of the yellow dots and to respond by pressing the “same” or “different” button. The task was administered on a hospital laptop and run via E-Prime 3.0 ^42^. A series of practice blocks were used to teach task demands. In study NCT05662280, participants completed a 10-minute version of the task that only included 3, 5, or 7 dot options, for a total of 72 trials across two blocks. In study NCT05102864, participants completed a 14-minute version of the task that included 1, 3, 5, or 7 dot options, for a total of 96 trials (across two, 48 trial blocks). Participants took a brief, self-determined break in between the two blocks. For consistency across datasets, only trials with 3, 5, or 7 dots were analyzed.

### Source Localization

Brainstorm’s processing pipeline uploaded each subject’s MRI anatomy and raw epoched EEG data file into MATLAB. Before beginning the source estimation, the subject’s anatomy was normalized in MNI space using the maff8, an SPM mutual information algorithm that implements a 4×4 linear transformation. This step establishes spatial uniformity between each of our subject’s brains and those imaged for previously published studies, allowing for generalizable observations of brain activity in a standard coordinate system. The subject’s EEG file was then co-registered with their MNI-normalized MRI anatomy. To co-register the EEG channels to electrode positions in this space, the ICMBM152 ANT Waveguard 64 EEG positions were added. Brainstorm converts the EEG coordinates from MNI to subject space to project the subject’s measured EEG activity onto the ICMBM152 ANT Waveguard 64 EEG channel positions.

MRI segmentation was applied to each subject’s data, generating boundary element method (BEM) surfaces of the scalp (1922 mm), outer skull (1922 mm), inner skull (1922 mm), and the skull (4 mm) using the Brainstorm BEM mesh generation method. Brainstorm’s OpenMEEGBEM forward modeling method calculate the gain matrix and computed a 3-layer BEM head model comprising of the scalp, skull, and brain with conductivities of 1, 0.0125, and 1 S/m respectively. The minimum norm imaging method factors current density map measurements with a constrained dipole orientation source model. Scouts were applied to the source model to extract the source activation of selected regions of interest exclusively. The Brainstorm processing pipeline bypasses the step of adding an MNI parcellation because Brainnetome atlases are registered with each subject’s MRI before being uploaded to Brainstorm. The Brainnetome Atlas, openly available for download at https://atlas.brainnetome.org/download.html, is a computed parcellated brain map representing the in vivo connectivity of 210 cortical and 36 subcortical regions ^43^. The DLPFC was defined by the Brainnetome Atlas scouts of the ventral area of region 9/46, and the PPC was defined by the Brainnetome Atlas scouts of the rostral-dorsal area of region 39. The left and right regions were averaged for a single DLPFC and a single PPC. The scout time series for these regions of interest were then extracted as a .mat file for later analysis.

### EEG Recording & Preprocessing

Recording was conducted at a 2000Hz or 2048 Hz sampling rate with CPZ set as reference electrode. Data was resampled to 250 Hz, line noise was removed (using EEGLAB *Zapline*), then high-pass filtered at 1 Hz and low-pass filtered at 80 Hz. EEGLAB’s *clean_rawdata* function was used to remove channels when a) signal was flat for 5+ seconds, b) high-frequency line noise was >4 standard deviations (SDs) above the mean, and c) when there was < .80 correlation with nearby channels. This function also performed artifact subspace reconstruction bad burst rejection for data > 20 SDs outside the mean and removed bad data periods with > 25% of channels out of acceptable range. Rejected channels were spherically interpolated and data was re-referenced to average. Independent component analysis (ICA) was conducted with the picard algorithm. Using EEGLAB’s *ICLabel*, components with a .8 or higher likelihood of being eye or muscle artifact were rejected. An epoch was rejected if it contained a value outside the ±150uV range.

